# Pyronaridine Tetraphosphate Efficacy Against Ebola Virus Infection in Guinea Pig

**DOI:** 10.1101/2020.03.20.001081

**Authors:** Thomas R. Lane, Christopher Massey, Jason E. Comer, Alexander N. Freiberg, Huanying Zhou, Julie Dyall, Michael R. Holbrook, Manu Anantpadma, Robert A. Davey, Peter B. Madrid, Sean Ekins

**Author notes:** To whom correspondence should be addressed: Sean Ekins, E-mail address. Boston University, National Emerging Infectious Diseases Laboratories, 401P, 620 Albany Street, Boston, MA 02118. Co-first authors, identified alphabetically by SE.

## Abstract

The recent outbreaks of the Ebola virus (EBOV) in Africa have brought global visibility to the shortage of available therapeutic options to treat patients infected with this or closely related viruses. We have recently computationally identified three molecules which have all demonstrated statistically significant efficacy in the mouse model of infection with mouse adapted Ebola virus (ma-EBOV). One of these molecules is the antimalarial pyronaridine tetraphosphate (IC_50_ range of 0.82-1.30 µM against three strains of EBOV and IC_50_ range of 1.01-2.72 µM against two strains of Marburg virus (MARV)) which is an approved drug in the European Union and used in combination with artesunate. To date, no small molecule drugs have shown statistically significant efficacy in the guinea pig model of EBOV infection. Pharmacokinetics and range-finding studies in guinea pigs directed us to a single 300mg/kg or 600mg/kg oral dose of pyronaridine 1hr after infection. Pyronaridine resulted in statistically significant survival of 40% at 300mg/kg and protected from a lethal challenge with EBOV. In comparison, oral favipiravir (300 mg/kg dosed once a day) had 43.5 % survival. The *in vitro* metabolism and metabolite identification of pyronaridine and another of our EBOV active molecules, tilorone, which suggests significant species differences which may account for the efficacy or lack thereof, respectively in guinea pig. In summary, our studies with pyronaridine demonstrates its utility for repurposing as an antiviral against EBOV and MARV, providing justification for future testing in non-human primates.

**Importance:** There is currently no antiviral small molecule drug approved for treating Ebola Virus infection. We have previously used machine learning models to identify new uses for approved drugs and demonstrated their activity against the Ebola virus *in vitro* and *in vivo*. We now describe the pharmacokinetic properties of the antimalarial pyronaridine in the guinea pig. In addition, we show that this drug is effective against multiple strains of EBOV and MARV *in vitro* and in the guinea pig model of Ebola virus infection. These combined efforts indicate the need to further test this molecule in larger animal efficacy studies prior to clinical use in humans. These findings also may be useful for repurposing this drug for use against other viruses in future.

## Introduction

Repurposing drugs for different diseases offers the opportunity to take a molecule that is approved for one clinical use and apply it to another disease, potentially accelerating its application and approval (1–3). There have been many articles on this approach and its successes (4). Several examples demonstrate repurposing compounds for the Ebola virus (EBOV) (5–7) which is a member of the virus family *Filoviridae* and pathogenic in both humans and non-human primates, causing severe hemorrhagic fevers (8) with mortality rates as high as 90% (9, 10). The recent outbreaks of EBOV in Africa have highlighted the need for new antiviral drugs for this and other emerging viruses to counter the human and financial cost (11, 12). The outbreak in Western Africa in 2014-2016 killed over 11,000 and caused over $53bn in economic damage (13). The current ongoing outbreak in the Democratic Republic of the Congo, in which well over 2200 people have died to date at the time of writing, and where the current case fatality is ∼67% (14)), emphasizes this need for new drugs while there is still no FDA approved drug for this disease. Several small molecule drugs such as favipiravir (15, 16) and most recently remdesivir (17) have been tested against EBOV in patients, although it is unclear whether any of them demonstrate efficacy (18, 19). We have previously used a computational approach with a published high-throughput screen of 868 molecules tested in a viral pseudotype entry assay and an EBOV replication assay (20, 21). This computational model enabled us to virtually screen several thousand compounds and identify three active compounds: tilorone, quinacrine and pyronaridine (22). All of these molecules inhibited EBOV in HeLa cells but not Vero cells, and they all demonstrated significant *in vivo* activity in the mouse-adapted EBOV (ma-EBOV) efficacy model (23–25). Pyronaridine (EC_50_ range of 420 nM-1.14 µ M (22, 26)) has been previously described in detail (27). It is the major component of the EU-approved antimalarial Pyramax, which is a combination antimalarial therapy with artesunate and pyronaridine and is approved for this use in the Democratic Republic of the Congo as well as other countries (e.g. South Korea). Our recent assessment of pyronaridine treated ma-EBOV-infected mice in range-finding studies indicated that a single 75 mg/kg i.p. dose which when given 1hr after infection resulted in 100% survival and statistically significantly reduced viremia on study day 3 (25). Additional studies in ma-EBOV-infected mice demonstrated that we could dose pyronaridine (75 mg/kg) 2 or 24hrs post-exposure without affecting survival (25). This was mirrored in our previous tilorone EBOV mouse study with treatment doses at 30 mg/kg q.d. (23). The pyronaridine mouse efficacy study also provided preliminary insights into how pyronaridine may possess antiviral activity as cytokine and chemokine panels suggested immunomodulatory actions during an EBOV infection (25). Our recent follow-up studies with the structurally related quinacrine (24) indicated this and many other structurally related antimalarials are active against EBOV *in vitro* (25) and may have a similar mechanism of action as all are known or suspected to be lysosomotropic amines. Such lysosomotropic compounds can diffuse across the membranes of acidic cytoplasmic organelles in their unprotonated form, then become protonated in the acidic environment, causing substantial accumulation in these organelles (28), which has the potential to ultimately impact lysosomal function.

It should be pointed out that there are many FDA approved drugs for which the mechanism is unknown. It is only in recent years that we have started to unravel the mechanism of action of such drugs that were approved decades ago (29, 30). The focus of our current efforts is on assessing pyronaridine and other clinical stage compounds as possible treatments for EBOV. Our studies to date with tilorone, quinacrine and pyronaridine may also provide compounds which could be combined as EBOV therapies for future assessment. While the focus of this study is testing pyronaridine, tilorone was also evaluated alongside favipiravir in the guinea pig model of EBOV infection to assess whether the efficacy observed in mouse would also be observed in this species. *In vivo* studies in the guinea pig would if successful then lead the way for studies in non-human primates.

## Results

### Testing vs EBOV and MARV strains

Pyronaridine, tilorone, and quinacrine were all previously discovered using machine learning models for EBOV and tested against the Mayinga strain (22). We now demonstrate that they block the entry stage of infection in a pseudotype assay (Fig. S1). Even though EBOV and MARV are distantly related (31) we also now show these three compounds are active against MARV Musoke strain in HeLa Cells (Fig. S2). These compounds were found by two of our groups to be similarly efficacious against multiple EBOV (Kikwit, Mayinga and Makona) and MARV (Musoke and Angola) (Fig. 1 and Table 1) strains in HeLa cells.

**FIG 1.**
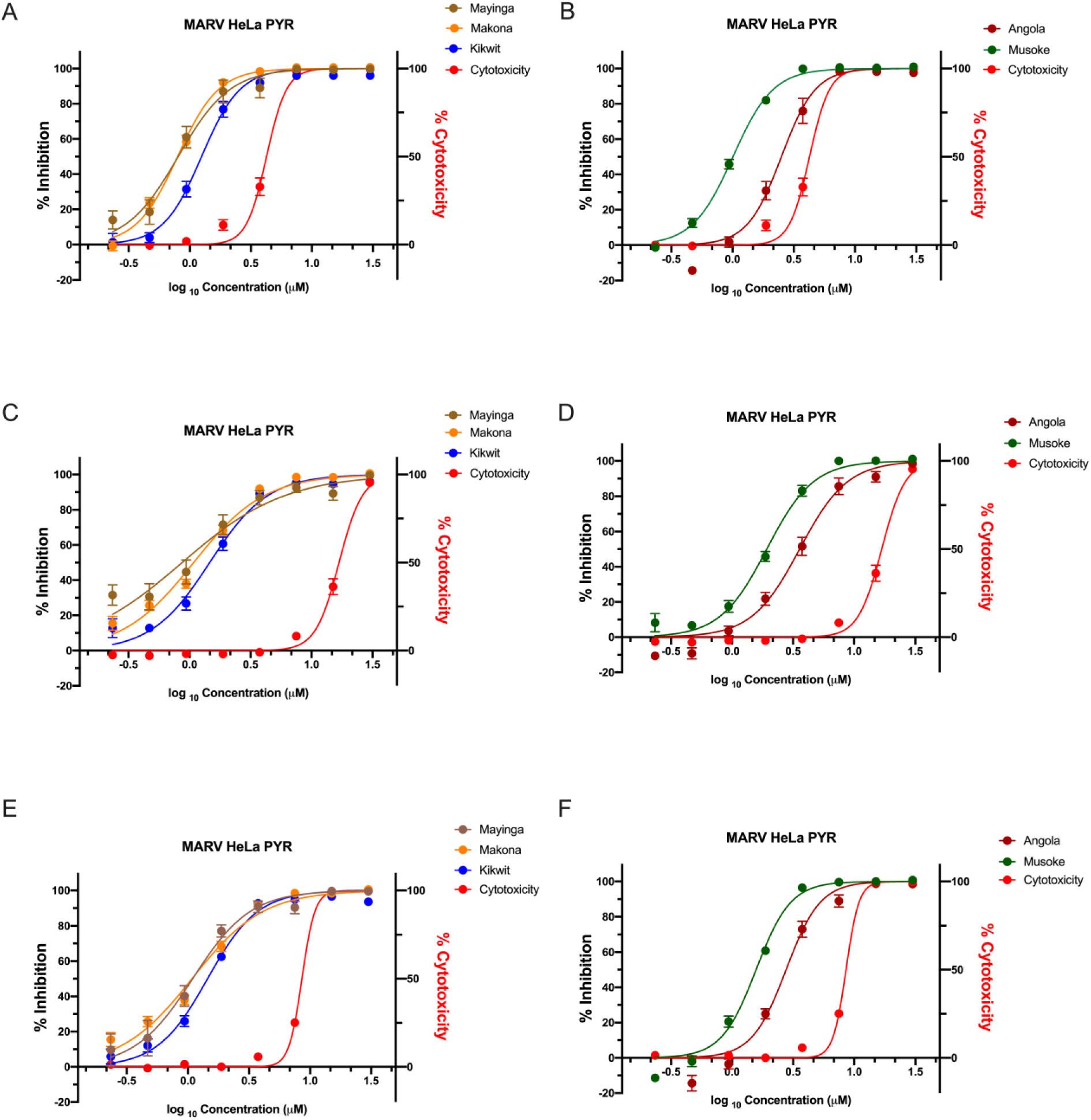
Pyronaridine, tilorone and quinacrine efficacy and cytotoxicity dose response relationship against multiple strains of EBOV (Kikwit, Mayinga and Makona) and MARV (Musoke and Angola) in HeLa cells. (EBOV/Kik, Mak, May: MOI 0.21; MARV/Ang: MOI 021; MARV/Mus: MOI 0.4).

**Table 1.**
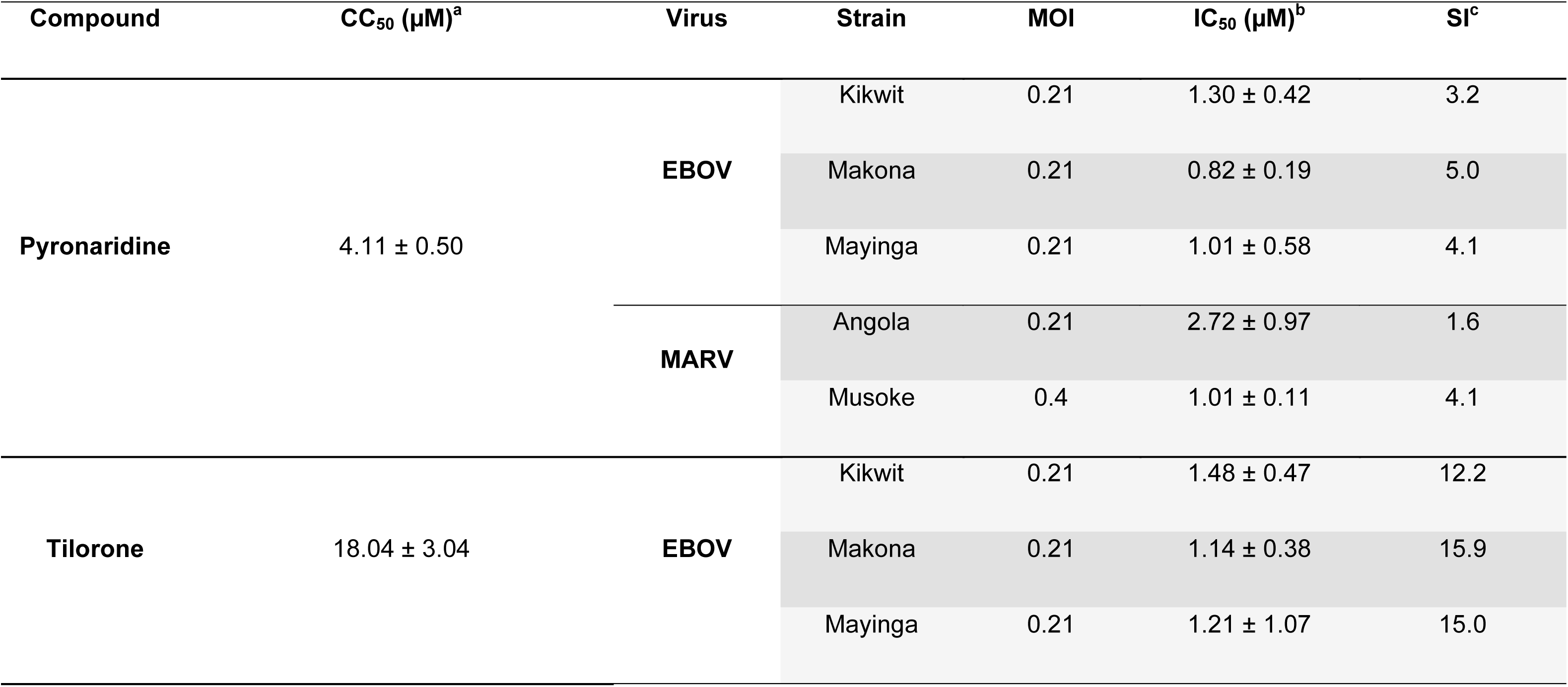
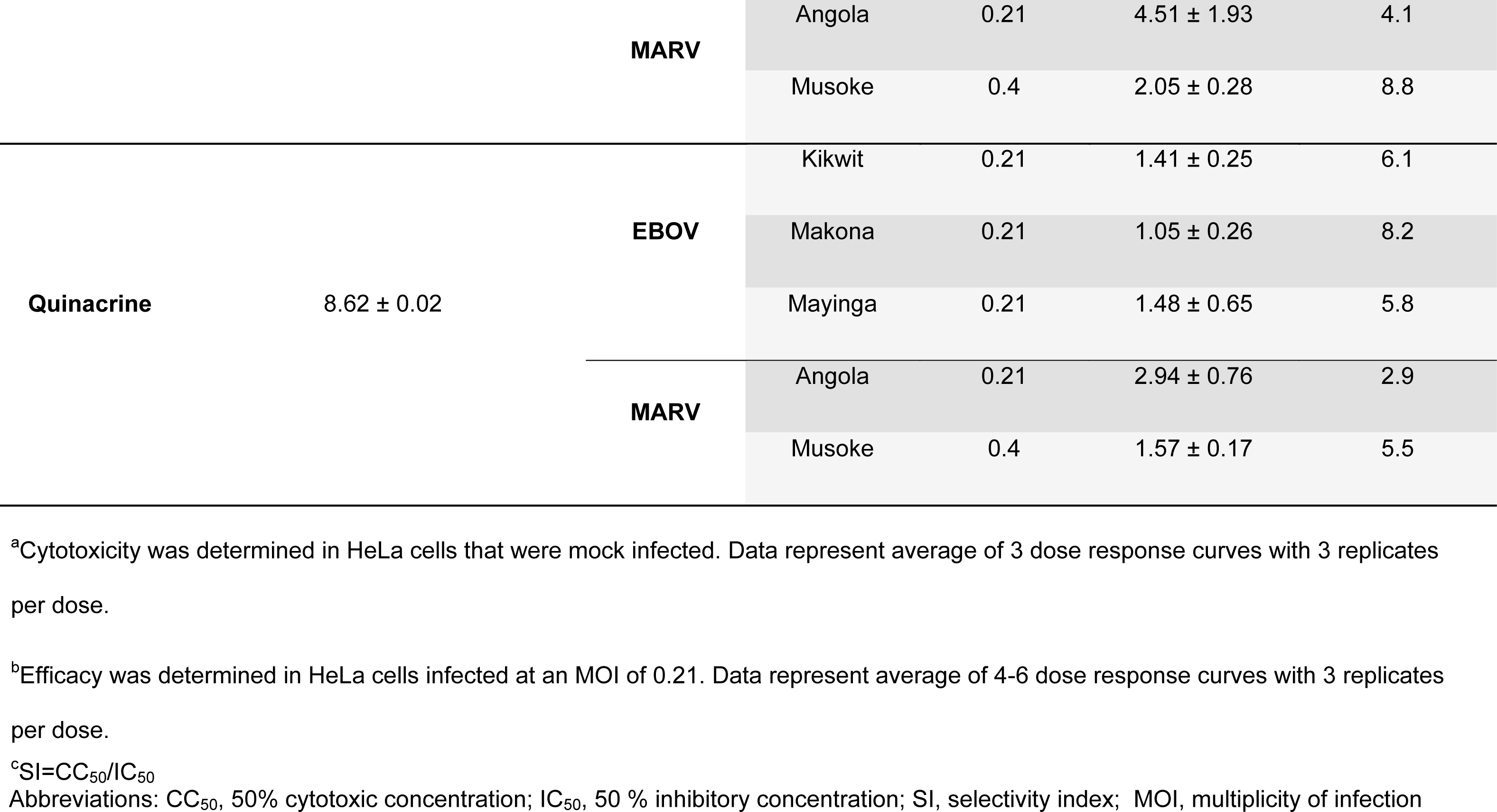
Pyronaridine, tilorone and quinacrine (± SD) show a similar efficacy against multiple strains of EBOV (Kikwit, Mayinga and Makona) and MARV (Musoke and Angola) in HeLa cells. Analysis via a F-test rejects the hypothesis that the CC_50_ and the respective IC_50_ are the same for each of the compounds evaluated (EBOV, Mayinga, tilorone is ambiguous).

### Metabolic stability across species

We have previously characterized the *in vitro* metabolic stability of pyronaridine in mouse, guinea pig, non-human primate and human (25). We have now performed a comparison for tilorone, quinacrine and chloroquine (a known lysosomotropic compound (32)) under similar conditions. Pyronaridine liver microsome (LM) metabolic stability increased in the order of guinea pig, non-human primate, human and then mouse. Tilorone had a similar species-LM stability relationship, with an increase in the order of guinea pig, non-human primate, mouse, followed by human. Chloroquine differed, with LM metabolic stability in the order of mouse, non-human primate, guinea pig and then human. Finally, quinacrine metabolic stability increased in the order of non-human primate, mouse, guinea pig and then human (Table 2). The CYP2D6 substrate probe dextromethorphan metabolism closely paralleled the species differences observed for pyronaridine and was also used to normalize the t_1/2_ (Table S1 and S2).

**Table 2.**
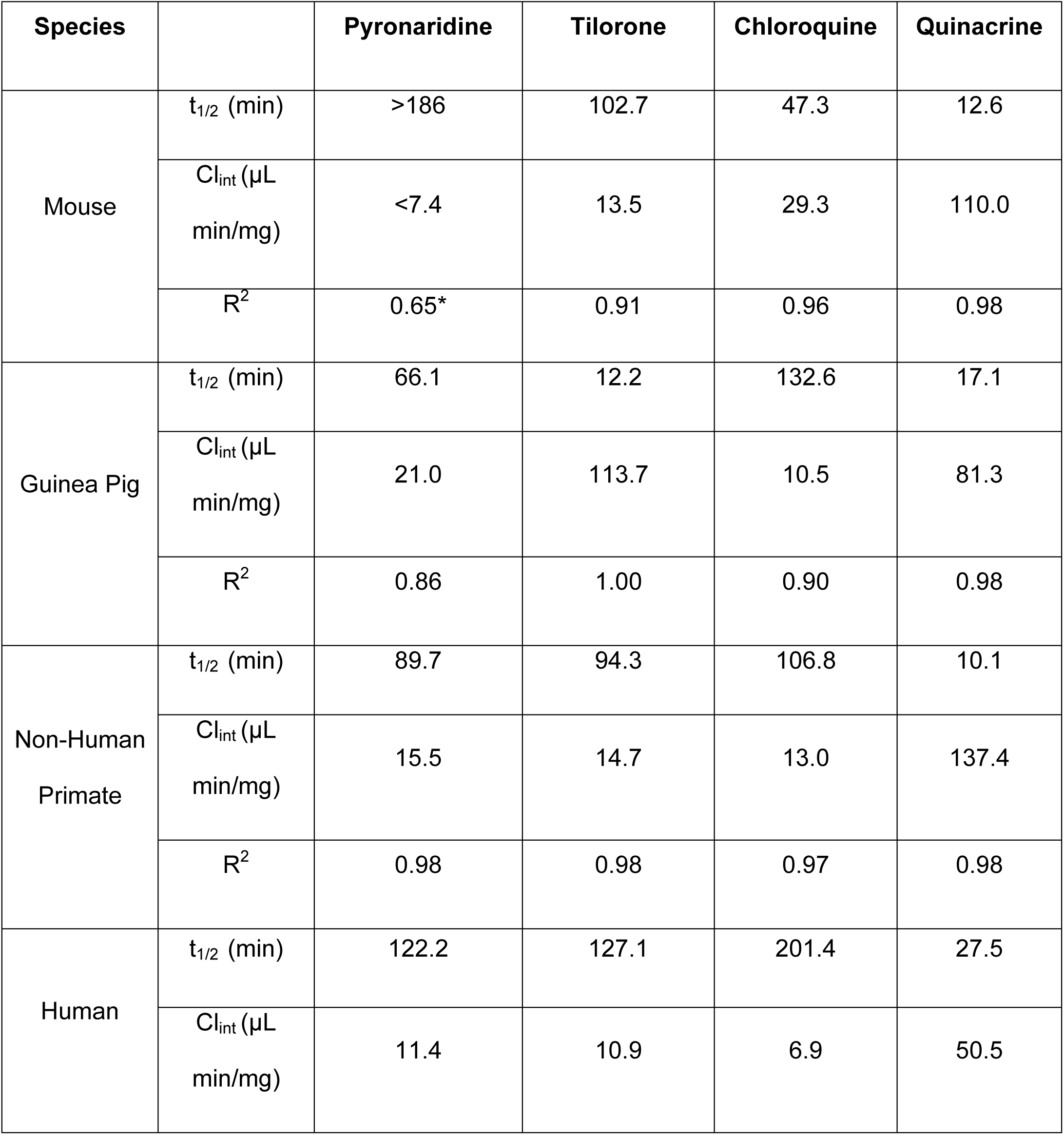
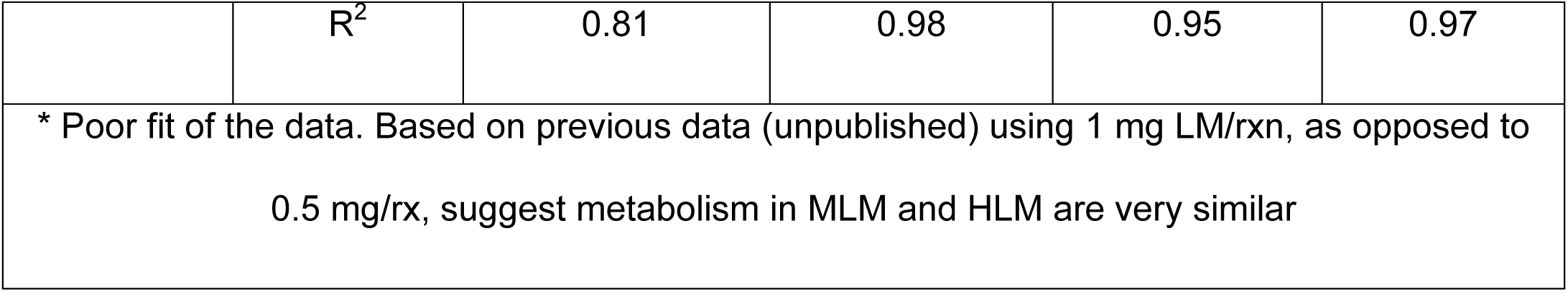
Liver microsomal metabolic stability across species

### Metabolite identification across species

We have previously characterized the pyronaridine metabolites produced in mouse microsomes (25). We have now evaluated the metabolites of multiple compounds with *in vitro* activity against EBOV (pyronaridine, tilorone, quinacrine and chloroquine) across multiple species (human and guinea pig) (Fig. S3-S6). The relative peak area abundance (%) for pyronaridine mono-oxygenation was much higher in guinea pig as compared to human liver microsomes. Tilorone *N*-deethylation and mono-oxygenation was higher in guinea pig relative to both mouse and human. Quinacrine O-demethylation was also 2-3 times higher in guinea pig. In contrast, chloroquine mono-oxygenation was highest in mouse relative to other species. Overall, guinea pig metabolism for these compounds in LMs differed substantially as compared to the other species tested.

### Guinea pig dose range-finding toxicity

The maximum tolerated dose of pyronaridine was evaluated in Hartley guinea pigs (Fig. 2). In the pyronaridine i.p.-dosed groups the highest dose level of 300 mg/kg was acutely toxic, with 4 of 6 guinea pigs found dead within 30 mins post injection. In addition, one died on day 7 and surprisingly the final surviving guinea pig showed no abnormal clinical observations. For the 200 mg/kg i.p.-dosed guinea pigs, 2 of 6 were found dead on days 5 and 6 and one met criteria for euthanasia on day 6. The remaining surviving guinea pigs from this group were found prostate on day 7 (Fig. 2D). No abnormal clinical observations were noted for guinea pigs administered either 125 mg/kg pyronaridine or vehicle via i.p. administration for the duration of the study. Oral dosing however had drastically reduced toxicity, with only 1 of 6 having any abnormal clinical observations at 600 mg/kg, which was detected directly following administration. This animal was found breathing rapidly for 6 mins, but fully recovered 2 hr post dose. There were no abnormal clinical observations at 300 or 125 mg/kg via oral administration. Based on these results, the maximum tolerated dose (MTD) for a single pyronaridine dose was determined as 125 and >600 mg/kg for i.p. and oral administration, respectively (Fig. 2A). Additionally, the maximum tolerated dose of tilorone was also tested in guinea pigs (Supplemental Methods, Supplemental results, Fig. S7A).

**FIG 2.**
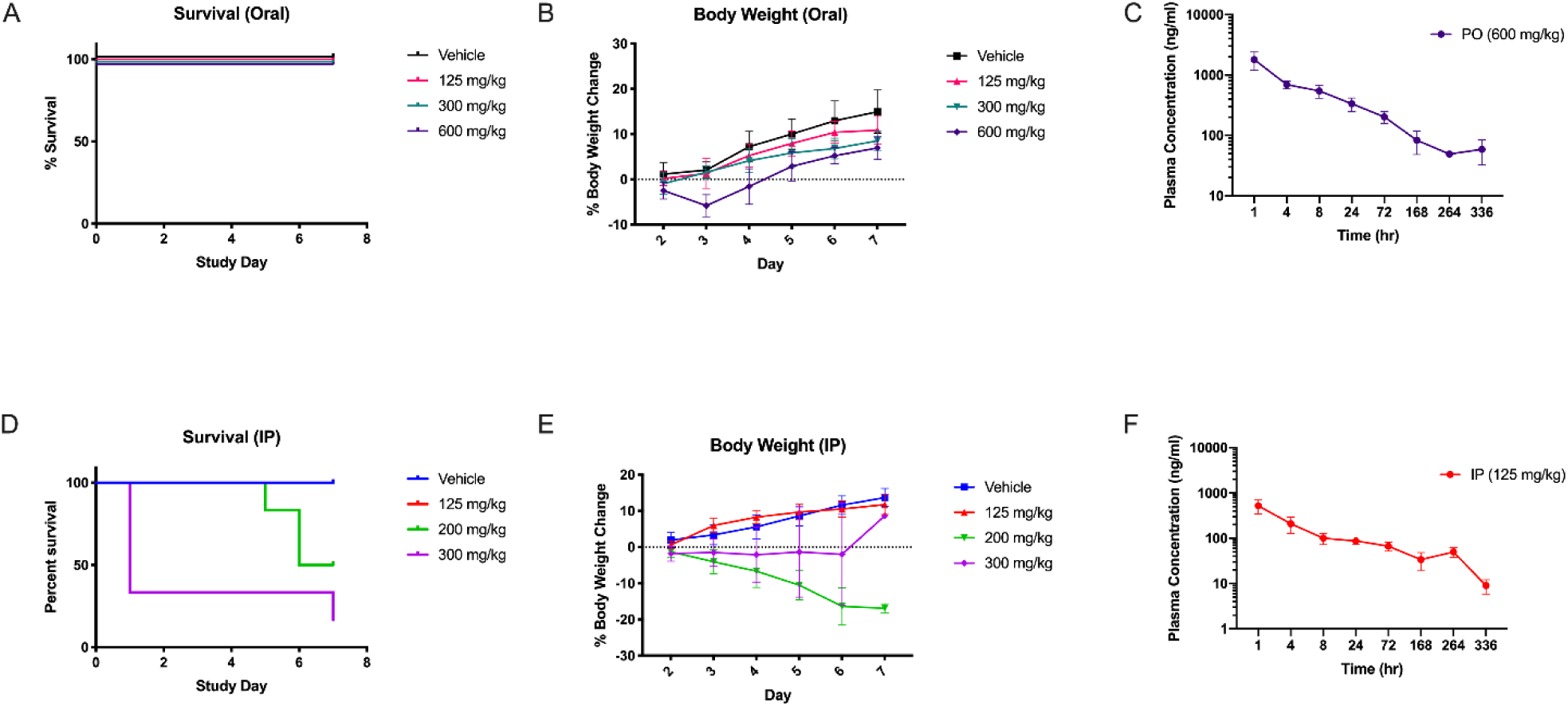
Guinea pig dose range-finding toxicity and Pharmacokinetics profile of Pyronaridine administered via oral gavage (A,B,C) or by intraperitoneal injection (D,E,F)

### Guinea pig pharmacokinetics evaluation of pyronaridine

The pharmacokinetics of pyronaridine was evaluated in Hartley guinea pigs (Fig. 2C, F). After an initial rapid absorption phase, the pyronaridine plasma profile exhibited a distribution phase at about 1hr, then a prolonged phase with plasma drug concentrations remaining essentially unchanged, or slightly higher until about 72hrs. All samples for animals dosed orally and i.p. contained measurable levels of pyronaridine though 336 and 168 hrs, respectively (LLOQ = 1 ng/ml). The plasma drug levels were analyzed using noncompartmental modeling allowing for the calculation of pharmacokinetic parameters (Table 3). Pyronaridine plasma levels reached the peak in the first sample, taken at 1 hr post administration. The elimination-phase t_1/2_ was calculated as 72.7 and 90.5 hrs for i.p. and oral administration, respectively. This is considerably shorter than the t_1/2_ found in humans and mice of between 195-251 hrs (33, 34) and 146 hrs (25), respectively. Maximum concentration of unbound drug in plasma (C_max_), area under the concentration-time curve from time zero to the last measurable concentration (AUC_last_), and area under the concentration-time curve from time zero to infinity (AUC_inf_) are provided in Table 3. Additionally, the pharmacokinetics of tilorone was also evaluated in guinea pigs (Supplemental Methods, Supplemental results, Fig. S7B, C and S8).

**Table 3.** Mean pharmacokinetics data in male guinea pigs treated with pyronaridine

|  |  |  |  |  |  | **C<sub>max</sub>**<br>**(ng/ml)** |  | **AUC<sub>last</sub>**<br>**(hr\*ng/ml)** |  | **AUC<sub>inf</sub>**<br>**(hr\*ng/ml)** |  |
| --- | --- | --- | --- | --- | --- | --- | --- | --- | --- | --- | --- |
| **Dose**<br>**(mg/kg)** | **Administration** | **Sex**<br>**(n=3)** | **T<sub>1/2</sub>**<br>**(h)** | **SE** | **T<sub>max</sub>**<br>**(h)** | **Mean** | **SE** | **Mean** | **SE** | **Mean** | **SE** |
| 125 | IP | M | 72.7 | 9.3 | 1 | 523 | 175 | 16,565 | 5269 | 17,430 | 5428 |
| 600 | Oral | M | 90.5 | 3.9 | 1 | 1800 | 348 | 50,964 | 4406 | 58,783 | 6712 |

### Pyronaridine efficacy and clinical observations

The efficacy of pyronaridine was evaluated in Hartley guinea pigs challenged with guinea pig adapted EBOV (gpa-EBOV). All animals in vehicle treatment (Group 1) succumbed to disease by study day 12 (100% mortality). Group 2 (pyronaridine 300 mg/kg) and Group 3 (pyronaridine 600 mg/kg) resulted in 40% and 33% survival, respectively. Among the 16 animals in the combined Group 4 (favipiravir 300 mg/kg), 7 GPs survived through the end of the 21-day study period; the other 9 guinea pigs succumbed to disease by study day 14 (43.75% survival, Fig. 3A). It should be noted that an animal in Group 2 was euthanized due to a clinical score of 4 on study day 2. Ebola disease in guinea pigs does not progress this rapidly so it is unlikely the animal succumbed to disease and it is more likely the animal incurred esophageal trauma because of the oral gavage technique, therefore this animal has been removed from data analysis.

**FIG 3.**
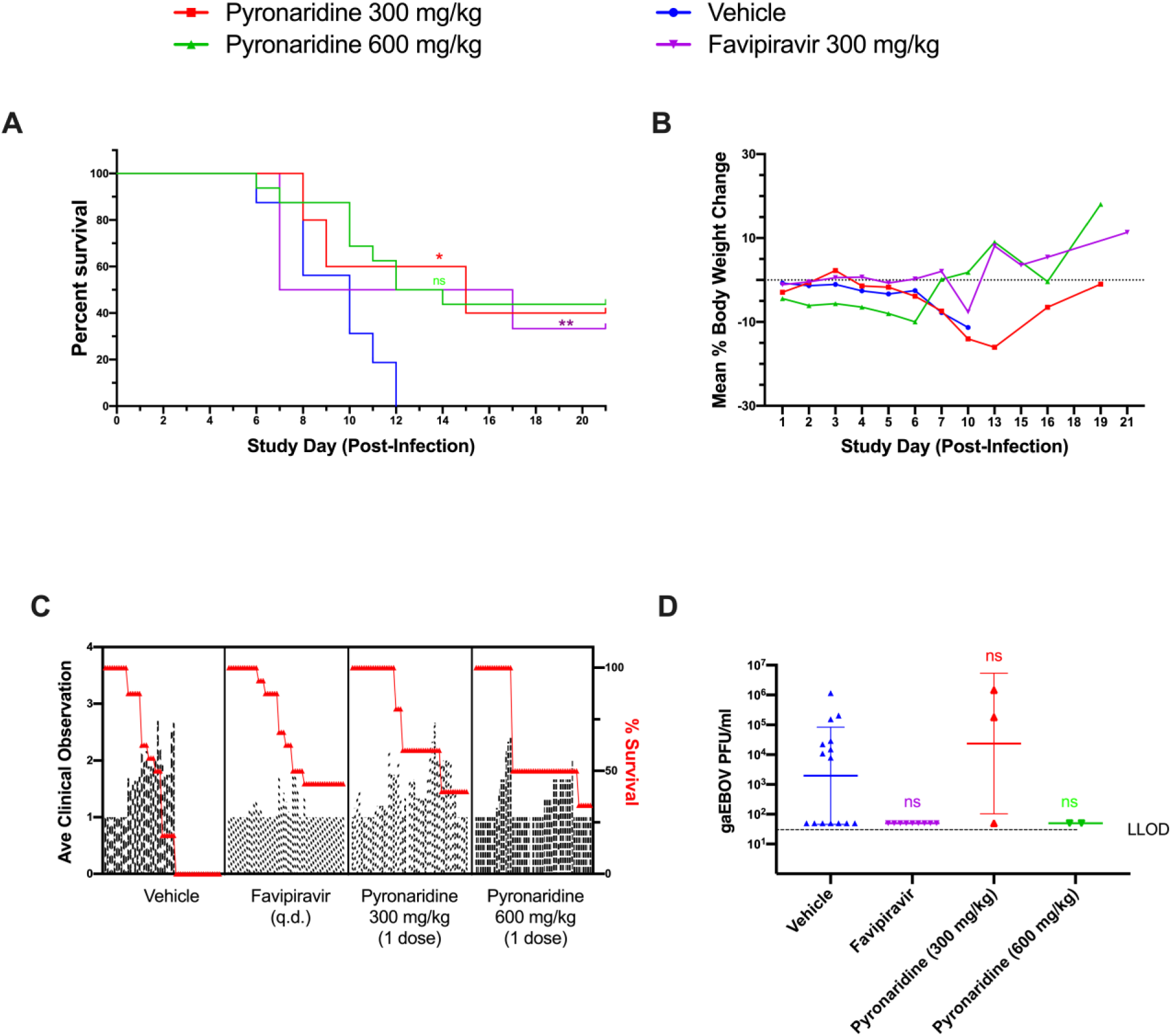
Guinea pig dose range-finding efficacy. gaEBOV efficacy data. Data from favipiravir and vehicle-treated (combined, n=16) groups were combined from our own two independent studies in order to strengthen their predictive power. (A) The survival curves between pyronaridine 300 and 600 mg/kg, favipiravir and vehicle. Asterisks represent significant difference from the vehicle (Log-rank (Mantel-Cox) test; Pyronaridine p=0.0307; Favipiravir p=0.0014). B. Mean percent body weight change from SD0 C. Mean clinical scoring results with overlaid percent survival. D. Plaque assay for viable EBOV in sera (GPs sacrificed based on clinical score) with Dunnett’s T3 multiple comparisons test. Statistical significance was calculated with log-transformed plaque assay data using a Dunnett’s T3 multiple comparisons test (Forsythe and Welch ANOVA) with the vehicle designated as the control. The difference from the vehicle was not found to be significant. For the plaque assay gaEBOV viral load had LLOD of 100 PFU/ml. Quantified values below these where set to 0.5 x LLOD. Bars and error-bars represents the geometric mean and geometric SD.

Body weight results for each treatment group are summarized in Fig. 3B. Animals in Group 1 maintained body weight through study day 7 but decreased by study day 10, when all animals succumbed to disease. Group 2 animals body weight continuously decreased through study day 7, with surviving animals returning to pre-challenge body weights by study day 10. Group 3 animals dropped body weight following challenge, but the survivors reverted to pre-challenge body weight by the end of the 21-day study. It should be noted that an animal in this group had developed clinical signs late and succumbed to infection on study day 17. Animals in Group 4 remained at a consistent body weight until study day 10 where the mean body weight began to decrease briefly, but the surviving animals rebounded to pre-challenge weight by study day 13.

Plaque assay results from serum samples taken from animals that met euthanasia criteria during the scheduled observation times from all four groups are presented in Fig. 3D. Only 10 of the 20 terminal samples had detectable levels of viable gpa-EBOV: eight animals from Group 1 (4.87 x 10^7^ geometric mean PFU/mL) and two animals from Group 2 (8.15 x 10^5^ geometric mean PFU/mL). Interestingly, all the guinea pigs treated with favipiravir were below the limit of detection for the plaque assay (Fig. S9).

Clinical scoring results for each guinea pig was assessed on a scale ranging from 1-4, where a score of 1 indicated a healthy animal and a score of 4 was indicative of a moribund animal. All animals in each group succumbed at least partially to disease by study day 8 (Fig. 3A). In each group the mean clinical observation increased inversely with survival, with each pyronaridine-treated group having two independent increases in average clinical observations. Interestingly, during post-study day 8 the average clinical score in the favipiravir group seemed to be independent of survival (Fig. 3C).

### Favipiravir and Pyronaridine *in vivo* efficacy against EBOV

It should be noted that in order to increase the statistical power of our study we combined the favipiravir, positive-control data from our two individual experiments (n=6, n=10) as these used comparable approaches and were performed by the same group. Due to the small group sizes the data for the negative, vehicle controls (n=6, n=10) were also combined with the noted caveat of a variation in frequency of administration.

Favipiravir has been shown to protect guinea pigs from adapted Sudan Virus (35) however in our study it protected ∼44% of the animals against gpa-EBOV, with deaths starting on study day 6 and continuing until study day 14 (Fig. 3A). These were not statistically different from the treatment with pyronaridine 300 and 600 mg/kg with 40% and 33% survival, respectively (Log-rank (Mantel-Cox) test). Tilorone at the doses assessed were either comparable or had significantly reduced survival rates as compared to the vehicle (Fig. S9A). In summary, there was a statistically significant difference for both pyronaridine (300 mg/kg) and favipiravir when compared to the control combined from our two studies (Fig. 3).

The results in guinea pig are surprising as previous studies in mouse have shown pyronaridine, tilorone and favipiravir all protect mice infected with ma-EBOV (24, 25, 36). Interestingly, virus was only detected in serum samples collected at the time of euthanasia from one and two Guinea pigs from the vehicle and pyronaridine 300 mg/kg, respectively, in the current pyronaridine study. This differed in our tilorone study (Supplemental Methods, Supplemental Results, Fig. S7-S9) where virus was recovered from all but one serum sample harvested from the guinea pigs in the vehicle and tilorone-treated groups (Fig. S9D). Virus was not detected in guinea pigs treated with favipiravir (both survivors and non-survivors) and was statistically significantly reduced from vehicle (Fig. 3D).

## Discussion

There have been very few small molecule drugs that have reached the clinic for testing against EBOV, including favipiravir (37), GS-5734 (remdesivir) (17) and galidesivir (17). Favipiravir has demonstrated 100% efficacy in the mouse model of EBOV (38, 39), 83% protection in the Interferon α/β and γ Double Knockout Mice mouse model (40), 17% (41) to 50% (42) survival in the cynomolgous macaque and increased survival and decreased viral load (43) or unclear efficacy in humans (15, 16). Once daily IV dosed remdesivir has demonstrated 100% survival in non-human primates only (44) and on this basis has been tested in humans during the current EBOV outbreak. Recently, a significant portion of data was described (499 individuals) from a clinical trial involving the investigation of multiple therapeutics against EBOV (NCT03719586) with ZMapp (a monoclonal antibody cocktail) (45)), remdesivir, MAb114 (a monoclonal antibody) (46)) and REGN-EB3 (monoclonal antibody combination) (47)). These results showed that the antibodies REGN-EB3 and mAb114 had overall survival rates of 71% and 66%, respectively, and were much more effective with patients with low viremia levels. Both ZMapp and remdesivir were shown to be less effective with a 51% and 47% survival rates, respectively (19). Ebola generally has a wide variation in its fatality rates of between 25% to 90%, (average ∼50%). While these results are promising for the monoclonal antibodies, the delivery and administration of likely temperature sensitive treatments to remote areas in Africa is a potential issue. A highly stable small molecule drug that could be given orally as a single dose would be ideal and alleviate some of these logistical challenges that constitute the critical final stage of delivering a therapeutic to the patient.

From this present study, pyronaridine and several other drugs which we have identified have shown activity in several strains of EBOV and MARV *in vitro* (Fig. 1 and Table 1), indicating they may have a broad-spectrum activity against the virus family *Filoviridae.* Based on our pseudovirus data these would appear to be preventing entry of the virus (Fig. 1). Two of these drugs were studied further in the guinea pig model of EBOV infection. It is apparent that pyronaridine did not show as substantial of a difference in the survival rate in guinea pig as was observed for mouse (100% survival) (25). This may be because the half-life for pyronaridine is much shorter in the guinea pig, so efficacious plasma levels of drug are likely not maintained long enough (90 hrs). The pharmacokinetics varies in other species, where the half-life in mice and humans is approximately 140 hrs and 200 hrs, respectively. Many other small molecules have failed to progress beyond guinea pig for EBOV due to a lack of significant efficacy (7, 48–51). While antibodies have been successfully used in this animal model (52, 53), these failures may represent a significant limitation of the guinea pig model to extrapolate small-molecule efficacy against EBOV in humans. We have demonstrated there are substantial metabolic stability differences between mouse, non-human primate, human and guinea pig (25), with the latter having a considerably lower metabolic stability for pyronaridine (Table 2). This could be one explanation of why several drugs for EBOV perform well in mouse but fail in the guinea pig. While to our knowledge this has not been determined for favipiravir, it has been shown that the pharmacokinetics of this compound exhibit nonlinearity over dose and time in non-human primates (54), making these interspecies’ comparisons potentially much more complex. Based on this *in vitro* data it would also suggest the metabolic stability of pyronaridine in the non-human primate may also be poor, requiring a dose adjustment to retain efficacy in this model. Antibody therapeutics for EBOV are not likely to be metabolized by the same drug metabolizing enzymes; therefore, they may show more universal efficacy across species. We have also evaluated the metabolism of several drugs of interest in liver microsomes of various species under similar conditions (Table 2). The structurally unrelated tilorone also appears to have limited metabolic stability in guinea pig and is much more stable in non-human primate and human liver microsomes. Chloroquine and quinacrine show different patterns across species. Our comparison of metabolic stability with the substrate probe dextromethorphan may also point to the role of CYP2D family in the metabolism of pyronaridine. It has previously been shown that pyronaridine inhibits known substrates of CYP2D6 both *in vitro* (25) and *in vivo* (55), also suggesting that it may be a CYP2D6 substrate as well. We have supplemented these data with detailed metabolite identification for each of these four compounds available for the first time (Figure S3-6). It is unclear what effect EBOV infection would have on the metabolic enzymes such as the P450’s in the guinea pig. To our knowledge, favipiravir has not previously been tested orally against EBOV in guinea pig (though it has demonstrated survival rates of 83-100% in Sudan virus-infected guinea pigs (35)), and in this study we now demonstrate efficacy on a par with what was observed in non-human primates (41, 42). In comparison, survival after pyronaridine (300 and 600 mg/kg) treatment was not significantly different from oral-administered favipiravir in the guinea pig model ((Log-rank (Mantel-Cox) test), suggesting a similar efficacy. There was a statistically significant difference for both pyronaridine (300 mg/kg) and favipiravir when compared to the control combined from our two studies (Fig. 3). Our initial dose ranging work showed significant toxicity with pyronaridine when dosed i.p. in guinea pig (accumulation in the abdominal cavity) hence the focus on oral administration in this study. It should be noted that we used only a single dose of pyronaridine for all our efficacy studies, and it is feasible that more frequent dosing to overcome the lower half life may result in a higher exposure and subsequent increased survival rate.

Developing small molecule drugs for EBOV is extremely challenging. While high throughput screens have readily identified many FDA approved drugs as well as other candidate molecules with *in vitro* inhibitory activities against EBOV (20, 56), when these compounds are tested in guinea pig, to date, all of them have failed. For example, chloroquine (7, 48), azithromycin (7), amiodarone (49), iminosugars (50), BGB324 (51), NCK8 (51) and 17-DMAG (51) were all inactive in the guinea pig *in vivo* model. Interestingly, tilorone (57), chloroquine (32), azithromycin (58), amiodarone (59), BGB324 (60) and 17-DMAG (61) are all known lysosomotropic compounds and NCK8 is most likely as well (62). This may indicate that these compounds could have also failed due to a common antiviral mechanism that does not transcend species. In addition, there is recent evidence that the type-I IFN antiviral immune response in guinea pig is significantly different than in mouse or non-human primate (63), therefore guinea pig may not be an appropriate model to universally predict the antiviral response in humans. This is particularly relevant to EBOV, since viral susceptibility and adaptation to guinea pig was directly linked to differences in the immune response (64).

In total, these results for pyronaridine, tilorone and favipiravir may question the need for demonstrating efficacy against gpa-EBOV before expanding to the non-human primate model of Ebola virus infection. They also suggest that larger group sizes are required to show statistical significance and to allow for spontaneous animal deaths due to issues with oral gavage, which ultimately reduces animal group sizes.

In conclusion, the guinea pig *in vivo* data collected in this study points to ∼40% survival for pyronaridine and favipiravir against gpa-EBOV. The accumulated *in vitro* metabolic data indicates that the guinea pig may be a suboptimal model to predict the efficacy of these compounds to combat EBOV. This could be due to differences in EBOV mechanism and drug metabolism (e.g. species differences in the metabolic enzymes involved (65, 66)). Our combined *in vitro* and *in vivo* studies with pyronaridine demonstrate its potential utility for repurposing as an antiviral against different strains of EBOV and MARV. These efforts also provide ample justification for future testing of pyronaridine efficacy in non-human primates and possible provision of the combination drug Pyramax for future Ebola outbreaks.

## MATERIALS AND METHODS

### Ethics statement

All work with gpEBOV-challenged guinea pigs was approved by the University of Texas Medical Branch’s IACUC (IACUC protocol number 1805041 approved 5^th^ June 2018) and was done in accordance with all applicable sections of the Final Rules of the Animal Welfare Act regulations (9 CFR Parts 1, 2, and 3) and *Guide for the Care and Use of Laboratory Animals: Eighth Edition* (Institute of Laboratory Animal Resources, National Academies Press, 2011; the *Guide*). This work was conducted in UTMB’s AAALAC (Association for the Assessment and Accreditation of Laboratory Animal Care)-accredited GNL BSL4 laboratory.

### Chemicals and reagents

Pyronaridine tetraphosphate [4-[(7-Chloro-2-methoxybenzo[b][1,5]naphthyridin-10-yl)amino]-2,6-bis(1-pyrrolidinylmethyl)phenol phosphate (1:4)] (22) was purchased from BOC Sciences (Shirley NY). Favipiravir was purchased from AdooQ Bioscience (Irvine, CA). Tilorone and pyronaridine was purchased from BOC Sciences. Quinacrine and Chloroquine were purchased from Cayman Chemicals (Ann Arbor, Michigan) and Sigma Aldrich (St. Louis, MO), respectively.

### *In Vitro* ADME assays

*In vitro* ADME studies were performed by BioDuro (San Diego, CA).

### *In Vitro* liver microsome stability assays

The liver microsome solution (197.5 µL, 0.5 mg/ml protein concentration) (Sekisui Xenotech, Kansas City, KS) was aliquoted into 1.1 ml tubes, to which 2.5 µL of positive control and test compound stock solutions (100 µM in DMSO) were added. The tubes were vortexed gently, pre-incubated for 5 min at 37°C, then 50 µL of 5 mM NADPH or LM buffer (no NADPH buffer) was added into the tubes. For analysis, an aliquot of 30 µL was removed from each tube at 0, 5, 15, 30 and 60 min (without-NADPH reaction:0 and 60 min) and quenched with 300 µL of 5/10 ng/ml terfenadine/tolbutamide in methanol/acetonitrile (1:1, v/v). Samples were vigorously vortexed for 1 min and then centrifuged at 4,000 rpm for 15 min at 4°C. 100 µL of supernatant from each sample was transferred to tubes for LCMS analysis. The amount of parent compound was determined on the basis of the peak area ratio (compound area to IS area) for each time point (AB SCIEX 4500). Clearance rates were calculated by the equation: t1/2 = Ln(2)/ke and in vitro CL_int_ (µL/min/mg protein) = ke*Incubation volume/Microsomal protein amount, and ke using equation of 1st order kinetics:

### *In Vitro* Metabolite Identification of pyronaridine, quinacrine, chloroquine and tilorone in human, mouse, guinea pig liver microsomes

A DMSO solution of test compound was spiked into 50 mM KH_2_PO_4_ (pH 7.4) buffer containing liver microsome at a concentration of 1 mg/mL (Sekisui Xenotech, Kansas City, KS). The reaction was initiated by the addition of 1.0 mM NADPH to the reaction mixture. The final concentration of the test compound was 1 µM. After 0 min and 60 min incubation at 37°C, an aliquot was removed and the sample were precipitated with a 1:6 acetonitrile, quenching the reaction. The resulting mixture was centrifuged, and the resultant supernatants were dried at N_2_ stream, the resultant residue were reconstituted with 300 μL 10% acetonitrile/H_2_O (v/v) (0.1% FA) before LC-MS/MS analysis. The supernatant was used for LC-MS/MS analysis. All separations were performed on a ACQUITY UPLC BEH T3 1.8 µm column (2.1×100 mm) at 25°C with a flow rate of 0.3 mL/min. Mobile phase A consisted of 0.1% formic acid in water and mobile phase B consisted of 0.1% formic acid in acetonitrile. Chromatography used a step gradient by maintaining 1% mobile phase B for 5 min, 10% mobile phase B over 8.0 minutes, 20% mobile phase B over 2.0 min, 90% mobile phase B over 2 minutes, 95% mobile phase B over 2 minutes, then re-equilibration back to 1% B at 20 minutes. The total run time was 22 minutes. For all samples, a 5 μL aliquot of sample was injected. The mass spectrometer (HRMS, Q-Exactive Plus from Thermo Fisher) was operated in high-resolution, accurate-mass (HRAM) Orbitrap detection mode.

### Test article preparation for *In Vivo* studies

<u>Vehicle Preparation</u> (Pyronardine study): A solution of 20% Kolliphor HS 15 with Water for injection (WF)I was made to be used for the vehicle. Kolliphor HS 15 was melted at 60 °C. 10 ml of Kolliphor HS 15 was combined with WFI to a final solution volume of 50.0 ml (20% solution) and mixed using a vortex mixer for 30 seconds and then sonicated in an ultrasonic water bath for 25 minutes at 45°C. <u>Test Article Dose Preparation</u>: Dose formulations were prepared by mixing the pyronaridine in the vehicle to achieve the target concentration. The formulation was mixed by inversion 5-6 times and placed on an orbital shaker for 30±5 min. <u>Favipiravir Preparation</u>: A 0.5% solution of methylcellulose was prepared in sterile water. To this, the appropriate amount of Favipiravir was added, and the pH adjusted until the compound goes into solution. Favipiravir was prepared prior to challenge and stored at 4-8 °C.

### Guinea pig *in vivo* dose range-finding toxicity for pyronaridine

To assess the tolerance of pyronaridine and to select dose groups for pharmacokinetics studies, the drug was given to 5-6-week-old male and female Hartley guinea pigs (Vital River Laboratories) as a single dose by intraperitoneal (i.p.) administration or oral gavage (PO). The compound was formulated in 20% Kolliphor HS 15 (Solutol) in sterile water. There were 8 groups in total (i.p. and oral control groups), with 6 animals per group (3 male, 3 female). I.p. administration was 125, 200 and 300 mg/kg and oral was 125, 300 and 600 mg/kg, each with a dosing volume of 5 ml/kg. Clinical observations were initiated immediately post-dose and once daily up to 168 hrs post-dose.

### Guinea pig *in vivo* pharmacokinetics evaluation of pyronaridine

Guided by the dose range-finding study, the pharmacokinetics of pyronaridine in guinea pigs were initially assessed at 125 and 600 mg/kg (n=3; male) for i.p. and oral administration, respectively, concentrations at or below the MTD determined by the 7-day study. Pyronaridine for both oral and i.p. administration was solubilized in the same vehicle (20% Kolliphor HS 15). Blood was collected from the treated Guinea pigs at at 1, 4, 8, 24, 72, 168, 264 and 336 hrs post-dose for processing of plasma. All samples were analyzed, and drug levels were measured by liquid chromatography-tandem mass spectrometry (LC-MS/MS) with a lower limit of quantitation (LLOQ) of 1.0 ng/mL. Notably, in the pyronaridine i.p. dosed, 125 mg/ml group 2 of 3 Guinea pigs were found dead on days 14 and 17 post dose.

### Virus strains

For *in vivo* experiments, a well-characterized guinea pig-adapted Ebola virus stock (Ebola virus *Cavia porcellus*/ COD/1976/Mayinga-CDC-808012 (gpaEBOV)) was used for all efficacy studies (67). All work involving infectious gpa-EBOV was performed at the Galveston National Laboratory (GNL) biosafety level (BSL) 4 laboratory, registered with the Centers for Disease Control and Prevention Select Agent Program for the possession and use of biological select agents.

### Initial cell-based testing for inhibition against wild type MARV strain

MARV expressing GFP was used in testing against viral inhibition as outlined previously (31). In short, inhibitors were tested at 8 concentrations for activity. All treatments were done in duplicates, each replicate being on a different plate. Briefly, 4,000 HeLa cells (Ambion, Austin, TX) per well in 25 µl of medium were grown overnight in 384-well tissue culture plates. On the day of assay, test compounds were diluted to 200 µM concentration in complete medium. 25 µl of this mixture was added to the cells already containing 25 µl medium to achieve a concentration of 100 µM. 25 µl of medium was removed from the first wells and added to next well. This type of serial dilution was done 8 times to achieve concentrations of 100, 50, 25, 12.50, 6.25, 3.12, 1.56 and 0.78 µM. One hour after incubating with the compound 25 µl of infection mix containing wild type virus was used to infect cells. This resulted in a final concentration of 50, 25, 12.50, 6.25, 3.12, 1.56, 0.78 and 0.39 µM. Bafilomycin at final a concentration of 10 nM was used as a positive control drug. All virus infections were done in a BSL-4 lab to achieve a MOI of 0.075 to 0.15. Cells were incubated with virus for 24 hours. One day post infection cells were fixed by immersing the plates in formalin overnight at 4°C. Fixed plates were decontaminated and brought out of the BSL-4. Formalin from fixed plates was decanted and plates were washed thrice with PBS. MARV infected plates were immuno-stained using virus specific antibodies. Nuclei were stained using Hoechst at 1:50,000 dilutions. Plates were imaged and nuclei and infected cells were counted using Cell Profiler software.

Cells were permeabilized using 0.1% Triton X-100 (Sigma, Cat#T8787) in PBS and blocked for 1 h in 3.5% bovine serum albumin (Fisher-scientific-Cat#BP9704100), followed by immunostaining. Fixed cells were incubated with an anti-MARV VLP antibody (IBT bioservices, Cat#04-0005, 1:1500 dilution), overnight at 4°C. After 2 washes to remove any excess antibody cells were stained with anti-Rabbit Alexa-546 antibody (Life technologies, Cat#A11035). After 3 washes to remove any non-specific antibody nuclei were stained using Hoechst at 1:50,000 dilution and imaged on a Nikon Ti Eclipse automated microscope. Nuclei and infected cells were counted using CellProfiler software. Relative infection compared to untreated controls was plotted in GraphPad prism 8.2.1 software.

### Follow-up cell-based testing against EBOV and MARV strains

Compounds were tested in vitro against 3 strains of Ebola virus (Kikwit, Makona, Mayinga) and 2 strains of MARV (Angola, Musoke): Ebola virus/H.sapiens-tc/GIN/2014/Makona-C05 (EBOV/Mak, GenBank accession no. KX000398.1), Ebola virus/H.sapiens-tc/COD/1995/Kikwit-9510621 (EBOV/Kik, GenBank accession no. KU182905.1); Ebola virus/H.sapiens-tc/COD/1976/Yambuku-Mayinga (EBOV/May, GenBank accession no. KY425649.1); Marburg virus/H.sapiens-tc/AGO/2005/Ang-1379v (MARV/Ang, BioSample accession no. SAMN05916381); Marburg virus/H.sapiens-tc/KEN/1980/Mt. Elgon-Musoke (MARV/Mus, GenBank accession no. DQ217792). All virus stocks were propagated, and titers were determined by plaque assay on Vero E6 cells obtained from the American Type Culture Collection (Manassas, VA) as previously described (68).

The *in vitro* infection inhibition of the all the above filovirus strains was performed in HeLa cells. HeLa cells were seeded at 3 x 10^4^ cells/well in 96-well plates. After 24 hours (h), cells were treated with drugs at 2-fold dilutions starting from 30 µM. Cells were infected with virus 1 hr after the addition of the drugs in BSL4-containment at a multiplicity of infection (MOI) of 0.21 or 0.4. After 48 h, plates were fixed and virus was detected with a mouse antibody specific for EBOV VP40 protein (#B-MD04-BD07-AE11, made by US Army Medical Research Institute of Infectious Diseases, Frederick MD under Centers for Disease Control and Prevention contract) (68) or MARV VP40 protein (Cat# IBT0203-012, IBT Bioservices, Rockville, MD) followed by staining with anti-mouse IgG-peroxidase labeled antibody (KPL, Gaithersburg, MD, #074-1802). Luminescence was read on an Infinite® M1000 Pro plate reader (Tecan US, Morrisville, NC). The signal of treated, infected wells was normalized to uninfected control wells and measured (in percent) relative to untreated infected wells. Non-linear regression analysis was performed, and the 50% inhibitory concentrations (EC_50_s) were calculated from fitted curves (log [agonist] versus response [variable slope] with constraint to remain above 0%) (GraphPad Software, La Jolla, CA). Error bars of dose-response curves represent the standard deviation of three replicates. For quantitation of drug toxicity, HeLa cells were mock infected (no virus) and treated with drug dilutions under the same conditions as the infected cells. After 48 h, cell viability was measured using the CellTiter Glo Luminescent Cell Viability Assay kit according to manufacturer’s protocol (Promega, Madison, WI).

### VSV-EBOV-GP pseudotype virus assay

Vesicular Stomatitis Virus (VSV) pseudotyped with EBOV glycoprotein (GP) expressing a GFP reporter was generously provided by Dr. Wendy Maury (University of Iowa) and has been described previously (69, 70). VSV pseudotyped with EBOV glycoprotein was grown by infecting Vero cells (Ambion, Austin, TX) and then harvesting via filtration of the supernatant through 0.4 µM filters 24-30 hours after infection. Virus was then stored at −80 until use.

The cells were tested and imaged using the general methods outlined previously (31). In short, HeLa cells (Ambion, Austin, TX) were plated at a density of 20,000 cells/well of a 96 well plate. After attachment overnight, cells were pretreated with compounds for 1 hr at predetermined doses. The dosing series in this case was 25, 12.5, 6.25, 3.12, 1.56, 0.78, 0.39, 0.19, 0.09, 0.04, 0.02 and 0.01 µM. After 1 hr of incubation with compounds, the cells were infected with VSV pseudotyped with EBOV glycoprotein and expressing a GFP reporter. 24 hours after infection, cells were fixed in formalin. After fixation, formalin was washed off, nuclei stained with Hoechst and the cells imaged. Green cells (infected) and blue nuclei (total number of cells) were counted using cell profiler. Relative infection compared to untreated controls was plotted in GraphPad prism 8.2.1 software.

### *In Vivo* efficacy clinical observations and scoring

Twenty-four (24) experimentally naïve Hartley guinea pigs were assigned to four (4) gender balanced groups. Guinea pigs were anesthetized for dosing (challenge and treatment) via isoflurane inhalation. On study day 0 (SD0) all guinea pigs were challenged with 1000 PFU of gpa-EBOV in 0.2 mL of Minimum Essential Medium (MEM) via intraperitoneal (i.p.) injection. The viral dose administered was verified through plaque assay analysis of the prepared virus suspension.

Dosing for all pyronaridine and all tilorone groups occurred via oral gavage of test/control article on SD0 one hour (± 15 minutes) post-challenge. Favipiravir (300 mg/kg) was given by oral gavage once daily from SD0 through SD7. For the pyronaridine study on SD 3 and during unscheduled euthanasia blood was collected via retro-orbital bleed. For the tilorone study, blood was collected during scheduled and unscheduled euthanasia. Serum was harvested for viremia measurements via plaque assay.

Following challenge, animals were monitored daily by visual examination. Clinical scoring and health assessments were performed and documented at each observation using the scoring system wherein: 1= Healthy; 2= Lethargic and ruffled fur, 3= Sore of 2 + hunched posture an orbital tightening, 4 = Score of 3 + reluctance to move when stimulated, paralysis, unable to access feed and water normally or ≥ 20% body weight loss. Body weights were measured daily during the dosing period (SD0 – SD7) and then every third day until the study was completed. When animals reached a clinical score of 2, the frequency of clinical observations increased to twice daily, 4-6 hours after the initial observation. When the disease progressed, and the clinical score increased to a 3, the frequency of observations was increased to three times daily. All surviving animals were humanely euthanized on Study Day 21.

### Viral load determination

Serum was harvested from guinea pigs that met the euthanasia criteria. Serum harvested for plaque assay analysis was stored frozen (in an ultralow [i.e., −80°C] freezer) until the conclusion of the in-life portion of the animal study, after which samples were batch processed. For this assay, the limit of detection in this assay was 100 PFU/mL. For statistical analysis and graphing all values less than the LOD were assigned a value of one half the LOD.

## ACKNOWLEDGMENTS

We gratefully acknowledge the team at Bioduro for their considerable efforts on this project and in particular Mr. Dan Contoit for whom we dedicate this article to his memory. Dr. Joel S. Freundlich is kindly acknowledged for consultations. We acknowledge Elena Postnikova, Janie Liang, and Shuiqing Yu who performed the *in vitro* testing of compounds against multiple virus strains. Dr. Mupenzi Mumbere is kindly thanked for providing information on drug availability in the Democratic Republic of the Congo. H.Z., J.D., and M.R.H performed this work as employees of Battelle Memorial Institute (BMI). The findings and conclusions in this report do not necessarily reflect the views or policies of the US Department of Health and Human Services or of the institutions and companies affiliated with the authors.

